# Whole genome assembly and annotation of the lucerne weevil *Sitona discoideus*

**DOI:** 10.1101/2022.08.01.502324

**Authors:** Mandira Katuwal, Upendra R. Bhattarai, Craig B. Phillips, Neil J. Gemmell, Eddy Dowle

## Abstract

Weevils are a diverse insect group that includes many economically important invasive pest species. Despite their importance and diversity, only nine weevil genomes have been sequenced, representing a tiny fraction of this heterogeneous taxon. The genus *Sitona* consists of over 100 species, including *Sitona discoideus* (Coleoptera: Curculionidae: Entiminae), commonly known as lucerne (or alfalfa root) weevil. *Sitona discoideus* is an important pest of forage crops, particularly *Medicago* species. Using a dual sequencing approach with Oxford Nanopore MinION long-reads and 10x Genomics linked-read sequencing, we generated a high-quality hybrid genome assembly of *S. discoideus*. Benchmarks derived from evolutionarily informed expectations of gene content for near-universal single-copy orthologs comparison (BUSCO) scores are above 96% for single-copy orthologs derived from eukaryotes, arthropods, and insects. With a *de novo* repeat library, Repeatmasker annotated 81.45% of the genome as various repeat elements, of which 22.1% were unclassified. Using the MAKER2 pipeline, we annotated 10,008 protein-coding genes and 13,611 mRNAs. Furthermore, 68.84% of total predicted mRNAs and 67.90% of predicted proteins were functionally annotated to one or more of InterPro, gene ontology, and Pfam databases. This high-quality genome assembly and annotation will enable the development of critical novel genetic pest control technologies and act as an essential reference genome for broader population genetics and weevil comparative genetic studies.

## Introduction

Beetles (Coleoptera) are among the most diverse group of metazoans, with over 350,000 described species representing about one-fourth of all described species on the planet (Hunt, Bergsten et al. 2007, Stork, McBroom et al. 2015). Among beetles, the family Curculionidae (“true” weevils) contains over 60,000 described species, including many economically important invasive agricultural pests (Oberprieler, Marvaldi et al. 2007, McKenna, Sequeira et al. 2009). They present an excellent model for studying species diversity and evolution (Hunt, Bergsten et al. 2007) and insect–microbe association studies (Toju, Tanabe et al. 2013, Morera-Margarit, Pope et al. 2021). Despite their importance and striking diversity, only nine Curculionidae genomes are publicly available to date (Mei, Jing et al. 2022) limiting our understanding of this highly diversified taxon.

The lucerne weevil or alfalfa root weevil *S. discoideus* Gyllenhal 1834 (Coleoptera: Curculionidae: Entiminae) feeds mainly on *Medicago* species and occasionally on *Trifolium* species (Vink and Phillips 2007). They are strong flyers and are highly dispersive (Brockerhoff, Barratt et al. 2010). Although they are originally from southern Europe and northern Africa, they are currently found in many parts of the world, including Australia (Chadwick 1978), New Zealand (Esson 1975), the United States (O’Brien Charles 1982), Chile (Elgueta 1993), Argentina (Del Río, Lanteri et al. 2019), and South Africa (Geertsema and Volschenk 1993). *Sitona discoideus* populations in New Zealand are likely derived from a single introduction from Australia (Esson 1975) as both populations appeared closely related when compared to a Norfolk Island population (Vink and Phillips 2007). Because of its invasiveness, *S. discoideus* has established itself as a significant pest of lucerne in countries like New Zealand and Australia, costing millions of dollars annually (Hopkins 1982, Goldson and Muscroft-Taylor 1988). Adult *S. discoideus* feed on plant foliage, whereas larvae feed on the roots and root nodules. The latter stage is the more damaging as they can destroy the root nodules (Sue, Ferro et al. 1980) and significantly reduce plant productivity (Goldson, Dyson et al. 1985).

Like with many other pests, chemical control of *S. discoideus* is economically and ecologically unsustainable (Geertsema and Volschenk 1993) driving the preference for biological control strategies. Biological control has been a widely applicable tool for controlling invasive species worldwide as one of the most economical and long-term effective strategies (Clout and Williams 2009). However, heavy reliance on classical biological control of invasive species without integrating it into a more complete integrated pest management approach, increases the chances of failure because of imbalances caused by dramatic swings in pest populations (Street 2015). Hence, incorporating other control methods and information such as mechanical, cultural, ecological and genetic technologies is vital for the sustainability and effectiveness of biological control strategies (DiTomaso, Van Steenwyk et al. 2017). Furthermore, novel genetic tools possess a great potential to advance our understanding and enhance the precision and predictability of biological control (Goldson, Bourdôt et al. 2015, Street 2015).

Sequencing a pest species’ genome holds a myriad of opportunities, from the development of novel biocontrol strategies through genetic modification (Teem, Alphey et al. 2020) to comparative genomic studies to understand the underlying genetic traits of interest such as parasite or pesticide resistance (Chilana, Sharma et al. 2012). However, the lack of reference genomes for the genus *Sitona*, including *S. discoideus*, hinders our genetic understanding of its biology, ecology, and evolution. Therefore, creating a reference genome for this pernicious weevil will be an essential addition to the genomic resources available for weevils and the subfamily Entiminae. The rapid development of sequencing technologies like long-reads and linked-reads and assembly algorithms can be utilized to generate reference genomes with high quality and contiguity as they reflect gene content and genome structure (Whibley, Kelley et al. 2021). Furthermore, these technologies allow us to resolve haplotype issues, particularly for creating de novo assemblies of a heterozygous diploid organism (Zhang, Wu et al. 2020).

Here, utilizing the dual sequencing approach with 10x Genomics linked-reads and Oxford nanopore long-reads, we present high-quality genome assembly and annotation of *S. discoideus*.

## Materials and methods

### Weevil sampling and pre-processing

*Sitona discoideus* specimens were collected from three sites: Lincoln (−43.64230, 172.47090), Hindon (−45.68701, 170.22198), and Grassmere (−43.055663, 171.759499), across the South Island in New Zealand, because of their availability. The adult weevils were collected from mixed grass/legume paddocks containing lucerne (*Medicago sativa*) using a modified leaf blower. Collected weevils were identified, and snap-frozen in liquid nitrogen and stored individually in a tube at -80 ^°^C until further use. *Sitona discoideus* in New Zealand are parasitized by an introduced endoparasitoid wasp, *Microctonus aethiopoides*, so they were dissected under a dissection microscope to determine their parasitization status, and tissues from a single non-parasitized weevil were used for each nucleotide extraction. The weevils were washed with double-distilled water before dissection and were dissected using 1x PBS buffer (Goldson and Emberson 1981).

### High Molecular Weight (HMW) genomic DNA extraction

The high molecular weight genomic DNA from the tissue of adult weevils was extracted following the 10x Genomics recommended protocol for single insect DNA purification (https://support.10xgenomics.com/permalink/7HBJeZucc80CwkMAmA4oQ2). Extracted DNA was subjected to bead clean-up using AMPure XP beads before the library preparation. DNA was quantified in Qubit, and quality was checked in nanodrop and then stored at -20 °C. Only high-quality DNA was further used for library preparation.

#### 10X Genomics library preparation and sequencing

DNA was size selected to remove fragments shorter than 40kb using the Blue Pippin (Sage Science, USA). After size selection, 5.96 ng/μl of HMW DNA was used for Chromium 10x linked read (10x genomics, USA) library preparations following the manufacturer’s protocol at the Genetic Analysis Service (GAS), University of Otago (Dunedin, New Zealand). The library was sequenced to generate 2 × 151bp paired-end reads on the Nova-seq platform (Illumina) at Garvan Institute, Australia. The Nova-seq yielded only about 30x the coverage of the estimated genome size of *S. discoideus* compared to the recommended 56x for the standard assembly coverage required by the Supernova assembler. Therefore, we further sequenced the same libraries on a single lane of rapid flowcell in Hi-seq 2500 (Illumina) at Otago Genomics Facility (OGF), University of Otago, Dunedin, New Zealand.

#### Oxford MinION library preparation and sequencing

Before the nanopore library preparation, extracted DNA was sheared five times with a 26-gauge needle (Terumo, Japan). We prepared four long-read sequencing libraries using DNA from three males and a female adult. Libraries were prepared using a ligation sequencing kit (SQK-LSK109) (Oxford Nanopore Technologies, Oxford, UK) following the manufacturer’s protocol. The libraries thus prepared were individually loaded onto four R9 chemistry flowcells (FLO-MIN106) and sequenced for 72 hours or till pore exhaustion.

#### mRNA sequencing

We extracted total RNA from the different tissues and sexes of *S. discoideus* using a Directzol RNA MicroPrep kit (Zymo Research), using the on-filter DNAse treatment as per the manufacturer’s protocol. Samples for mRNA sequencing included different tissues (head, abdomen, and gonads) from adult males and females. We extracted total RNA from each individual and tissue type separately. The extracted total RNA was accessed for quantity and purity using Qubit 2.0 Fluorometer (Life Technologies, USA) and nanodrop; high-quality samples were stored at -80^°^C until further processing.

RNA integrity was evaluated using a Fragment Analyzer (Advanced Analytical Technologies Inc., USA) at the OGF. The report yielded RNA quality number (RQN) values that ranged from 5.7 to 8.4, where seven out of 12 samples produced an RQN value of 7 or above. However, the collapse of the 28S peak, a widespread phenomenon for RNA extracted from insects (Winnebeck, Millar et al. 2010), might be the reason behind lower RQN values; thus, we determined the quality via the trace instead of relying on the RQN value. After the quality control step, 12 RNA samples were used for Truseq stranded mRNA libraries preparation. A single equimolar RNA library pool was generated and a single lane of 2×150bp paired-end sequencing on the HiSeq 2500 V2 Rapid Sequencing flowcell was carried out in at the OGF.

#### Transcriptome assembly

mRNA-seq reads were quality filtered using Trimmomatic (v.0.39) with options: SLIDINGWINDOW:4:20 LEADING:5 TRAILING:5 MINLEN:36. The filtered reads were *de novo* assembled using Trinity (v.2.8.6) with all the default options.

#### Genome size estimation

We performed flow cytometry analysis using a single head of *S. discoideus* with two biological replicates at Flowjoanna (Palmerston North, NZ), following the standard procedures described in (Bhattarai, Katuwal et al. 2022). Rooster red blood cells (RRBC) obtained from a domestic rooster were used as a reference sample. The raw data of nuclei peaks were analyzed using Flowjo (BD BioSciences, USA) followed by the sample’s calculation of pg/nuclei.

#### Assembly strategies and bioinformatics pipeline

We sequenced and assembled the genome of the lucerne weevil *S. discoideus* at a total coverage depth of approximately ∼100x using linked and long-read strategies. We tried several assemblers and pipelines; however, the final pipeline optimized several criteria, including the BUSCO scores (Seppey, Manni et al. 2019) for gene completeness and reference-free metrics (total length, number of contigs/scaffold, number of N’s per 100kbp, N50 values and ortholog completeness). The assembly pipeline is described below (Figure 1).

**Figure 1:**
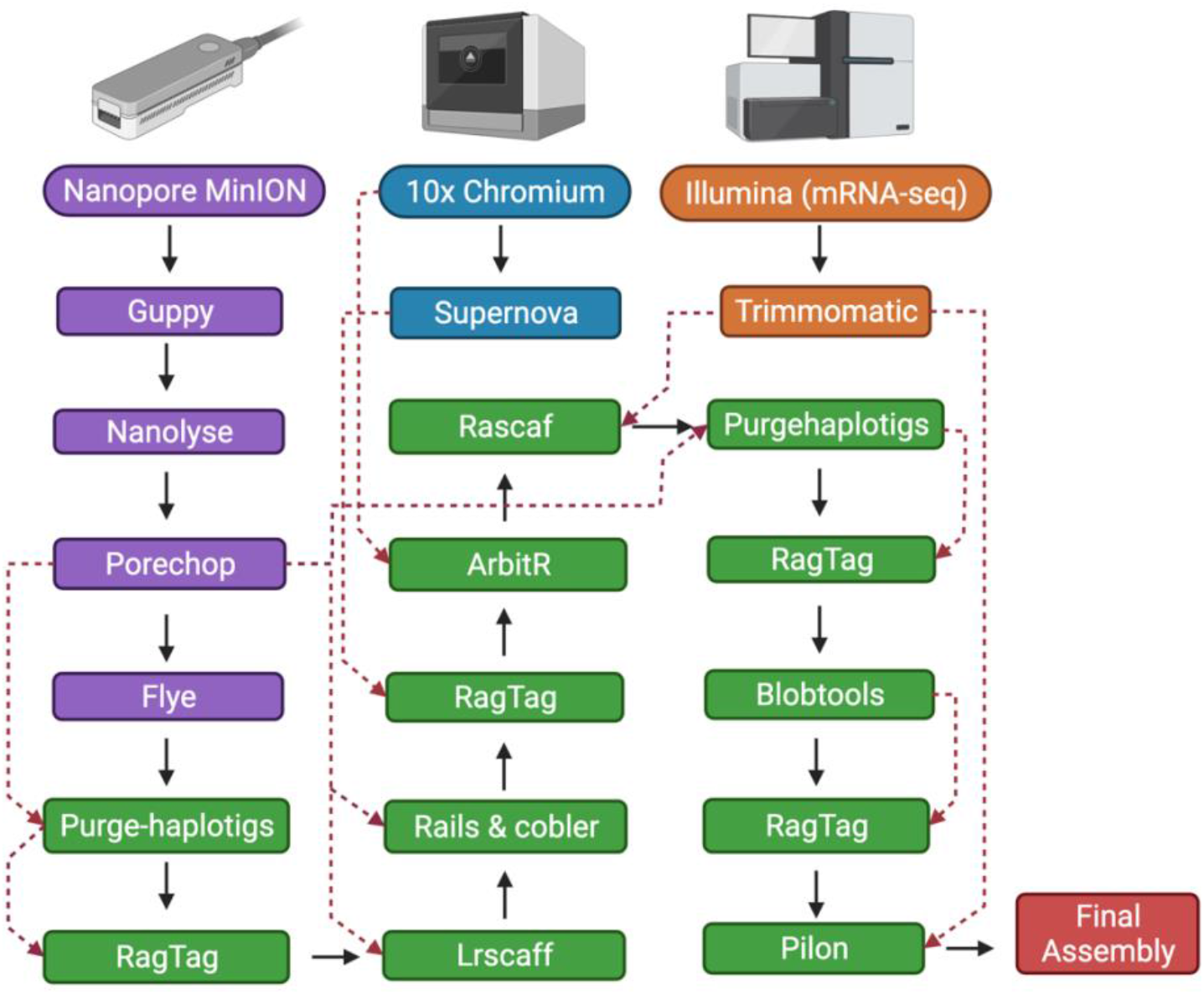
Schematic representation of the assembly pipeline for the *Sitona discoideus* genome. The black arrow represents the workflow, and the red dotted line represents the additional input data in the pipeline (Created with Biorender.com).

The sequencing reads from 10x Genomics Chromium linked reads sequencing were assembled using Supernova assembler v.2.1.1 with default parameters (Weisenfeld, Kumar et al. 2017). Multiple criteria, including gene completeness BUSCO scores (Seppey, Manni et al. 2019), Quast (Gurevich, Saveliev et al. 2013) and reference-free metrics (discussed above), were applied to assess the assembly quality. We used the “pseudohap” style of the Supernova “mkoutput” function to export the sequence in Fasta format.

The raw Nanopore reads from four MinION runs were base-called using Guppy v.5.0.7 (Oxford Nanopore Technologies) and adapter sequences were removed with Porechop v.0.2.4 (Wick, Judd et al. 2017). The filtered reads were assembled using Flye v.2.7.1 with default parameters (Kolmogorov, Yuan et al. 2019). The assembly statistics and the BUSCO percentage were better for the long-read Flye assembly than the linked-read Supernova assembly, so the Flye assembly was used as the primary assembly. Redundant and duplicated contigs were removed using Purgehalotigs (Roach, Schmidt et al. 2018).

Using the raw-filtered Nanopore reads as input, we scaffolded and gap-closed the purged assembly using Ragtag v.2.1.0 (Alonge, Soyk et al. 2019) Lrscaff v.1.1.11 (Qin, Wu et al. 2019), Rails v.1.5.1, and Cobbler v.0.6.1 (Warren 2016). The resultant assembly was again scaffolded with the Supernova assembly using Ragtag v.2.1.0 before being further scaffolded with ArbitR v.0.2 (Hiltunen, Ryberg et al. 2021) using the raw linked-read data. LongRanger (v.2.2.2) was used to align the linked-read data for ArbitR. We then used the mRNA-sequencing data to scaffold the genome further using Rascaf (Song, Shankar et al. 2016). Redundant and duplicated haplotigs of the genome were removed using the second round of Purgehaplotigs, with the discarded haplotigs being used for scaffolding the genome through Ragtag. The assembly was quality filtered to remove contaminants with Blobtools2 (Kumar, Jones et al. 2013, Laetsch and Blaxter 2017). We removed the contigs categorized as being from the bacterial and viral superkingdom, as well as contigs with less than five times coverage and less than 1000bp length. The reads discarded as shorter length (<1000bp) and low coverage (<5x) were used for the final scaffolding step via RagTag. The final assembly underwent two polishing rounds with Pilon (v1.24) (Walker, Abeel et al. 2014) using the Illumina mRNA-seq data.

#### Repeat content analysis

We generated a custom repeat library to aid with annotation for *S. discoideus* using multiple de novo repeat and homology-based identifiers, including LTRharvest (Ellinghaus, Kurtz et al. 2008), LTRdigest (Steinbiss, Willhoeft et al. 2009), RepeatModeler (Flynn, Hubley et al. 2020), TransposonPSI (Haas 2007) and SINEBase (Vassetzky and Kramerov 2013). We removed the redundant reads by concatenating the individual libraries and merging the sequences with over 80 % similarity using usearch v.11.0.667 (Edgar 2010) and then classified them with Repeat Classifier. We also mapped the sequences with unknown categories present in the library against the UniProtKB/Swiss-Prot database (e-value <1e-01), where the un-annotated repeat sequences were eliminated from the library. RepeatMasker v.4.1.2 (Chen 2004) was used with the final repeat library to produce a report for genome repeat content. Because of the time and the computational resources needed, we ran RepeatMasker with the quickest run option (−qq) and skipped the bacterial insertion element check option (−no_is). The repeat library was used to input the Maker2 (v.2.31.9) pipeline (Holt and Yandell 2011) during annotation.

#### Genome annotation

The weevil genome annotation was performed following the MAKER2 pipeline with three iterations, including both evidence-based and ab initio gene models. The evidence-based models were used for the first round of Maker, whereas the latter two rounds used ab initio gene model predictions. For the first round with the MAKER2 pipeline, 260,683 Mrna transcripts assembled through the Trinity pipeline (Grabherr, Haas et al. 2011), along with 5,281 mRNA and 13,621 Entiminae subfamily protein sequences downloaded from NCBI, were used as inputs. Snap and BUSCO trained Augustus was used for the latter two ab initio gene prediction rounds.

## Results and discussion

### Genome size estimates

The estimated genome size of *S. discoideus* from the flow cytometry was 946.215 ± 31.119 MB (mean ± SD). This is the first genome size estimation for the genus *Sitona*. This genome size estimation is within the range of those reported for other Curculionidae (162.6 to 2,025 MB) in InsectBase 2.0 (Mei, Jing et al. 2021).

### Transcriptome assembly

The Trinity pipeline produced an assembly of 205,961,878 bp length, with 260,683 contigs in total and 219 contigs with lengths more than 10,000 bp. The assembly has a GC ratio of 38.96% and N50 of 4512 bp. A BUSCO (v.5.2.2) analysis using the insecta_odb10 database found a complete BUSCO score of 96%.

#### Genome assembly

The 10x Genomics Chromium linked read library from a single individual weevil resulted in ∼311 million reads with coverage of ∼50x of the estimated genome size of *S. discoideus*. The Supernova assembler gave the assembly size of 340.11 MB, 144,095 total contigs with an N50 of 0.0068 MB and L50 of 6,063. We used Quast to estimate genome quality from the eukaryotic database, which reported a complete BUSCO of 35.97% and a partial BUSCO of 4.62% for the supernova assembly.

Similarly, sequencing with Nanopore MinION generated 30.4 Gb bases across ∼ 7.6 million reads. The N50 of the read length and the median read length were obtained from pycoQC (v 2.5.2) and were 13,500 and 1,230, respectively, with a median PHRED score of 13.23 (Supplementary Table 1). As the primary assembler, Flye generated an assembly length of 1,889 MB. The assembly resulted in 86,442 contigs with an N50 of 0.0785 MB and an L50 value of 6,634. A complete BUSCO score of 96.70% and partial BUSCO score of 1.32 % were reported with Quast using the eukaryotic database. The long-read assembly from Flye provided better contiguity and gene completeness (Table 1); therefore, we considered it our primary assembly to process further.

**Table 1.** Assembly statistics of *Sitona discoideus* genome assembly. Quast BUSCO scores are to its default Eukaryota database.

|  | Assembly length | No. of scaffolds | N50 | L50 | Ns per 100 kbp | BUSCO % (Quast) |  |
| --- | --- | --- | --- | --- | --- | --- | --- |
|  |  |  |  |  |  | Complete | Partial |
| Supernova assembly | 340,110,374 | 144,095 | 6,788 | 6,063 | 1,224.43 | 35.97 | 4.62 |
| Flye assembly | 1,889,302,767 | 86,442 | 78,533 | 6,634 | 1.56 | 96.70 | 1.32 |
| Final hybrid assembly | 1,172,662,393 | 6,835 | 297,589 | 952 | 3,338.46 | 96.04 | 1.32 |

Our *de novo* hybrid assembly resulted in a draft genome size of 1,172.66 MB spanning 6,835 contigs with N50 and L50 of 0.29 and 952 MB respectively, suggesting a contiguous assembly (Table 1). Furthermore, the assembly has 1,320 complete BUSCO genes using the insect database, representing 96.04% completeness, this includes 11.2% duplicated genes. The fragmented BUSCO was 1.32%. Similarly, BUSCO analysis against the Eukaryota database with Quast showed a complete and partial BUSCO of 96.04% and 1.32% respectively (Table 1). The contiguity and the completeness statistics shows that the assembly is of high quality (Figure 2). The assembly statistics after each round of processing are given in Supplementary Table 2.

**Figure 2.**
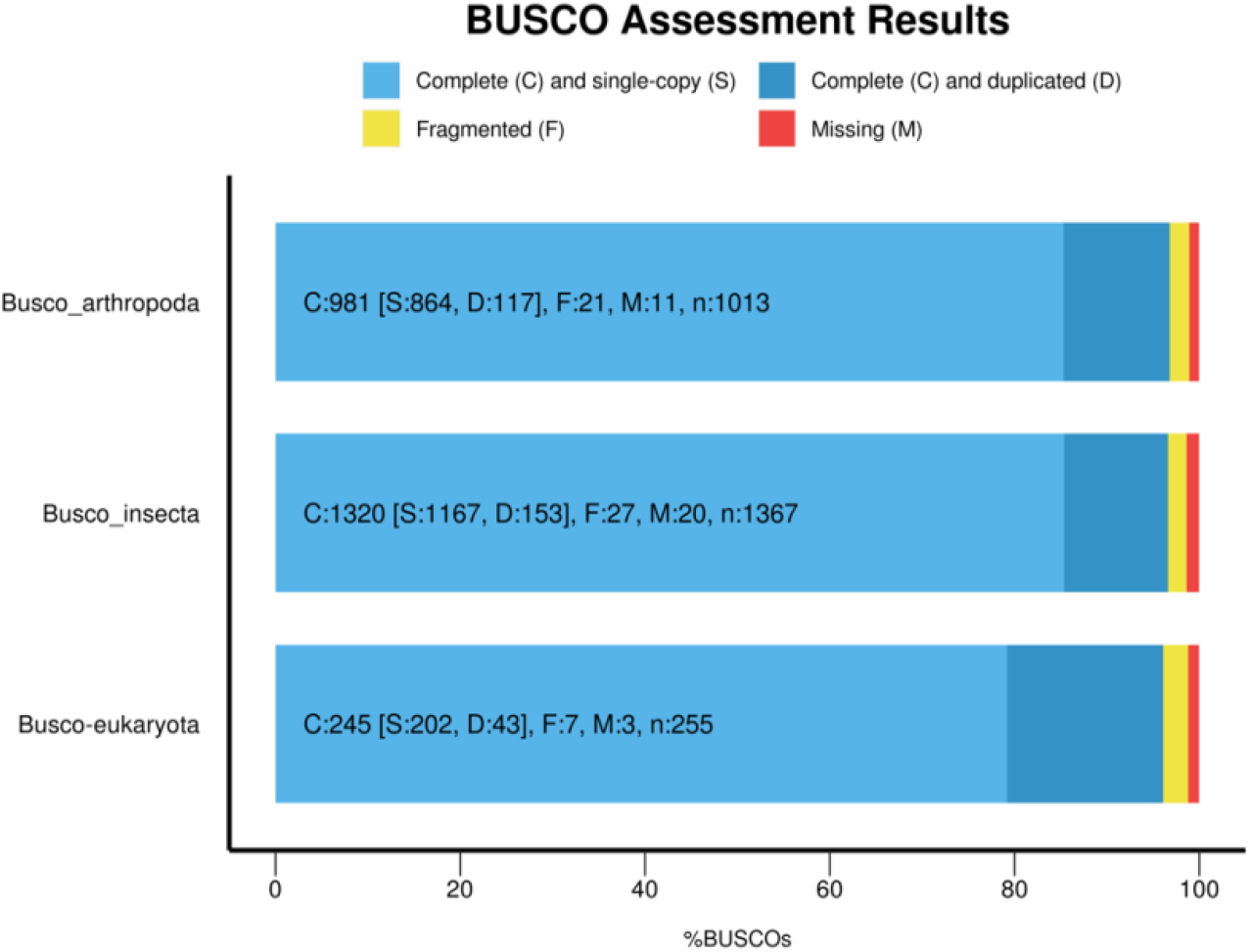
The BUSCO v.5 reports for the final hybrid assembly of the *Sitona discoideus* genome. BUSCO percentage (x-axis) from Arthropoda, Eukaryota, and Insecta (odt10) databases (y-axis) is shown in the bar plot. The light blue portion of the bar represents complete and single-copy orthologs, dark blue represents complete and duplicated orthologs, yellow represents fragmented BUSCO genes and red represents missing BUSCO genes.

As there are no publicly available genomes for the genus *Sitona*, we considered another forage pest weevil, the Argentine stem weevil (*Listronotus bonariensis*) to compare the genome with. The genome size of *L. bonariensis* is 1,112.4 MB with an N50 of 0.12 MB with a BUSCO completeness of 83.9% (Harrop, Le Lec et al. 2020). The *S. discoideus* genome size and N50 value are similar; however, the higher complete BUSCO score of 96.04% and a low partial BUSCO score of only 1.32% show that the assembly of *S. discoideus* is of high quality and contiguity in terms of its gene completeness.

#### Genome repeat contents

The Repeat Masker masked 81.45% of the genome as repeats. It reported 66.21% as the total interspersed repeats. This includes 25.06% as Retroelements, 19.95% as DNA transposons, 13.12% as Rolling-circles, and 21.2% as unclassified elements. Other repeat categories included Small RNA (0.35%), Satellites (0.36%), Simple repeats (1.37%), and Low complexity (0.04%) (Table 2).

**Table 2.**
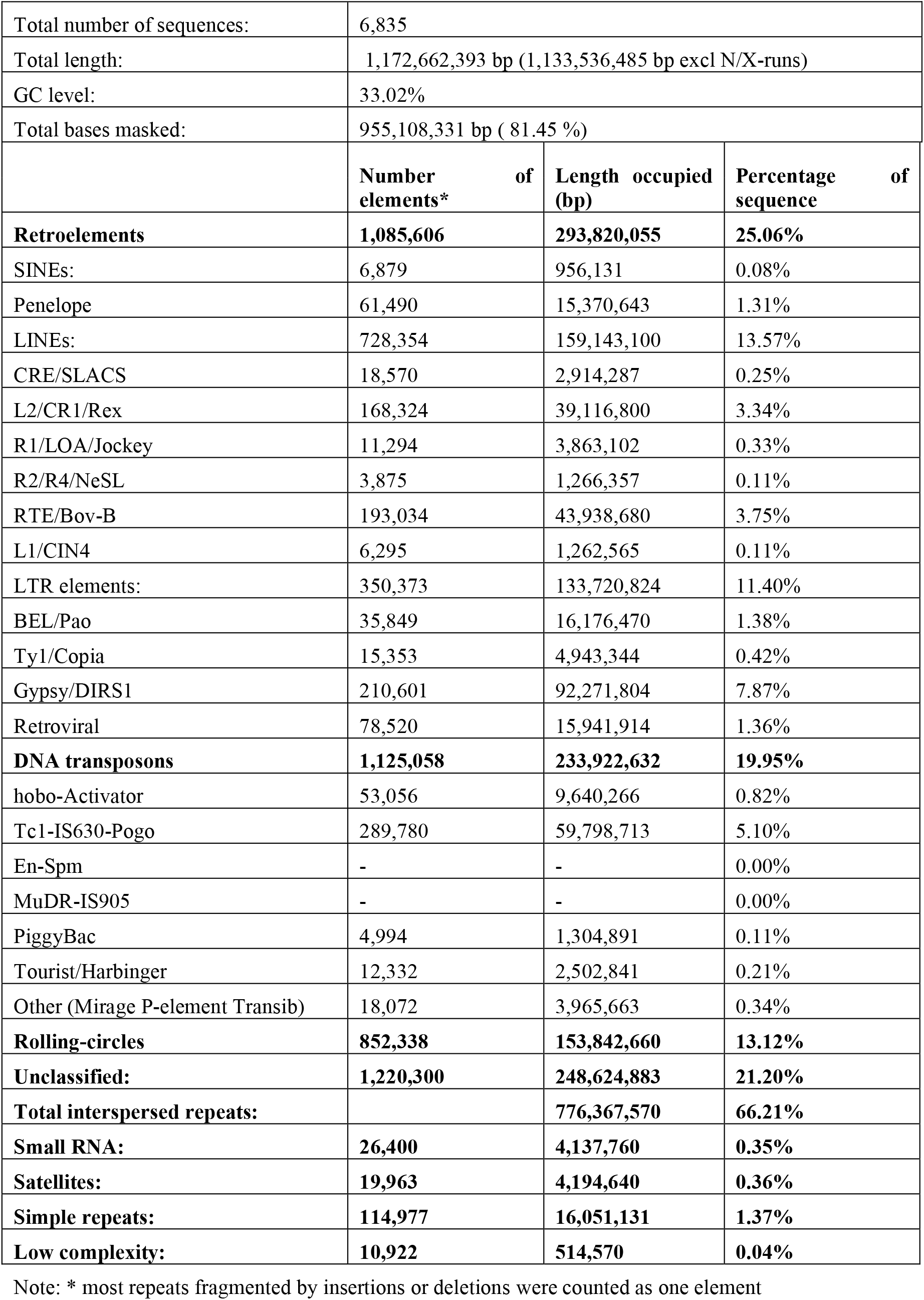
Repeat content analysis of *Sitona discoideus* genome

### Genome annotation

We identified 10,008 genes and 13,611 mRNAs in the assembled genome by combining evidence-based and ab initio gene models in the MAKER2 pipeline. The total gene length is 84.63 MB constituting 7.2% of the whole genome, and the mean gene length is 8,456 bp. Similarly, the longest gene annotated is 192,956 bp, and the longest CDS is 21,423 bp (Table 3). We also functionally annotated 68.84% of total predicted mRNAs and 67.90% of predicted proteins through either one or more of the InterPro, gene ontology, and Pfam databases (Supplementary Table 3). We got 64.1% and 62.2% of complete BUSCO scores for the annotated transcriptome and annotated proteins compared with the Insecta_odb10 database (Figure 3). We found that 99% of the gene models have an AED score of 0.6 or less, indicating highly confident gene prediction (Supplementary Figure 1). This assembly is comparable to a recently curated genome of the weevil *Pissodes strobi*, where 11,382 high confidence genes were reported, and 42.9% complete BUSCO genes were identified from Endopterygota_odb10 datasets (Gagalova, Whitehill et al. 2022). The number of annotated genes in *S. discoideus* is fewer than those reported in other coleopteran genomes, like the Easter egg weevil *(Pachyrhynchus sulphureomaculatus)* with 18,741 genes (Van Dam, Cabras et al. 2021), the pine beetle *Dendroctonus ponderosae* genome with 14,342 reported genes (Keeling, Yuen et al. 2013) and the Colorado potato beetle *Leptinotarsa decemlineata* genome with 16,533 genes (Schoville, Chen et al. 2018). A comparative study would shed more light on the difference in gene numbers between the beetles.

**Table 3.** Genome annotation summary for *Sitona discoideus*

|  |  |
| --- | --- |
| Total sequence length | 1,172,662,393 |
| Number of genes | 10,008 |
| Number of mRNAs | 13,611 |
| Number of exons | 88,311 |
| Number of introns | 74,700 |
| Number of CDS | 13,611 |
| Total gene length | 84,629,294 |
| Total mRNA length | 122,057,115 |
| Total exon length | 122,057,115 |
| Total intron length | 97,921,646 |
| Total CDS length | 16,937,688 |
| Shortest gene | 108 |
| Shortest mRNA | 108 |
| Shortest exon | 3 |
| Shortest intron | 5 |
| Shortest CDS | 18 |
| Longest gene | 192,956 |
| Longest mRNA | 192,956 |
| Longest exon | 11,865 |
| Longest intron | 162,917 |
| Longest CDS | 21,423 |
| mean gene length | 8,456 |
| mean mRNA length | 8,968 |
| mean exon length | 275 |
| mean intron length | 1,311 |
| mean CDS length | 1,244 |
| % of genome covered by genes | 7.2 |
| % of genome covered by CDS | 1.4 |
| mean mRNAs per gene | 1 |
| mean exons per mRNA | 6 |
| mean introns per mRNA | 5 |

**Figure 3.**
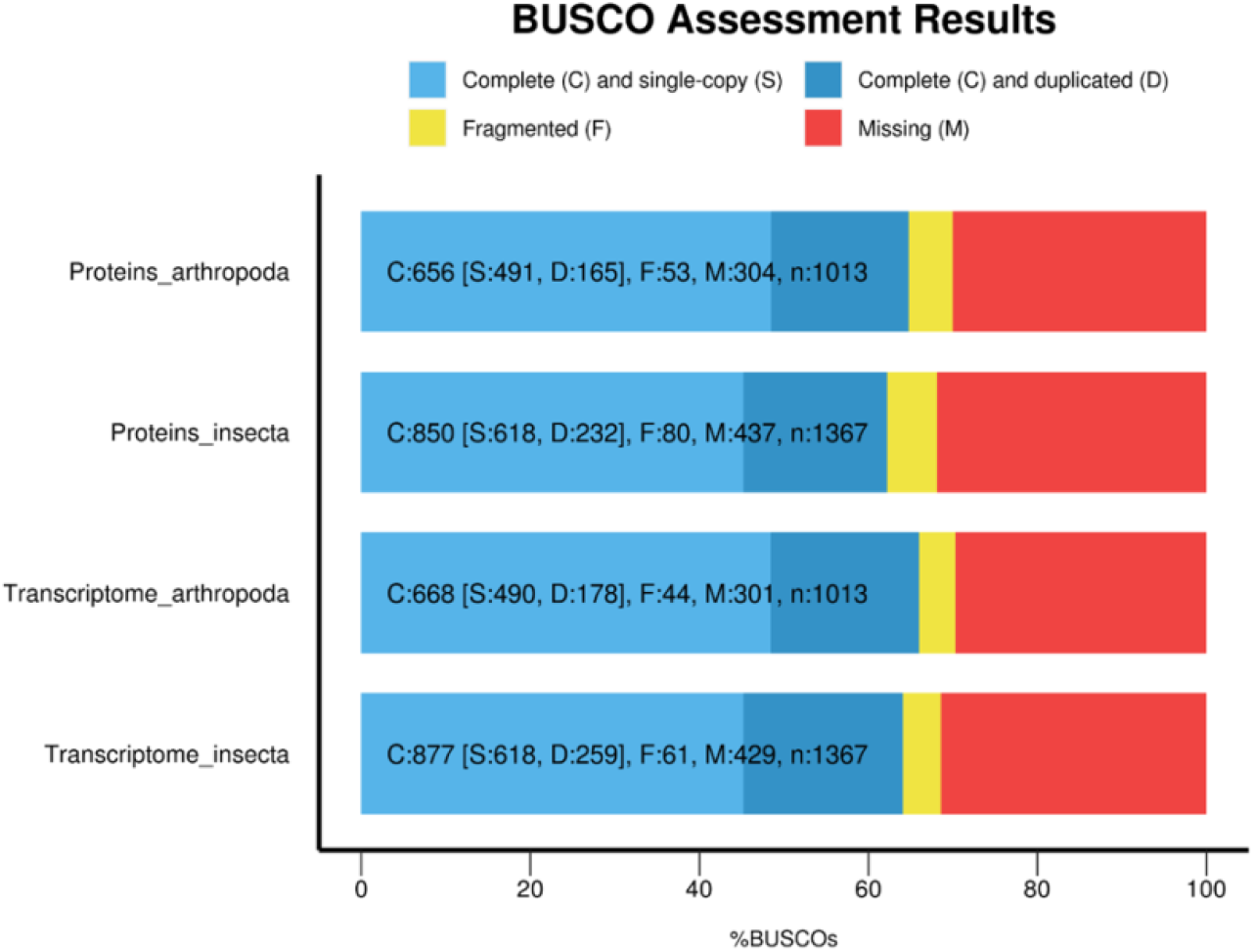
Annotation completeness through BUSCO database. The plot shows the BUSCO percentage (x-axis) for the annotated Proteins and Transcriptomes using Insecta_odb10 and Arthropda_odb10 database as indicated on the y-axis. BUSCO version 5 is used for the analysis.

Here, we report a high-quality assembled and annotated reference genome of *S. discoideus* using a dual sequencing approach, linked and long reads. This genome will aid in a wide range of genetic, genomic, and phylogenetic studies, particularly for the genus *Sitona* and other weevils of the subfamily Entiminae. More crucially this high-quality genome will guide our understanding of an economically important insect pest for which no management methods except biological control are available.

## Acknowledgments

We thank Dr. Scott Hardwick (AgResearch, Lincoln), Colin Ferguson (AgResearch, Invermay) for making the weevil samples available. We also thank the support team at New Zealand eScience Infrastructure (NeSI) for providing their HPC platform for the analysis.

