## Supplementary Table 1, Supplementary Table 2, Supplementary Table 3, Supplementary Figure 1 for "Whole genome assembly and annotation of the lucerne weevil *Sitona discoideus*"

**Supplementary Table 1. Nanopore sequencing output**

| Number of Reads | Total Bases | Median Read Length | N50 Length | Median PHRED score |
| --- | --- | --- | --- | --- |
| 5,272,527 | 22,461,690,000 | 1,230 | 13,500 | 13.23 |

**Supplementary Table 2. Assembly statistics after each round of processing**

| Assembly steps | Assembly length | No. of scaffolds | N50 | L50 | Ns per 100 kbp | Busco % (Quast) | |
| --- | --- | --- | --- | --- | --- | --- | --- |
|  |  |  |  |  |  | Complete | Partial |
| Supernova | 340,110,374 | 144,095 | 6,788 | 6,063 | 1,224.43 | 35.97 | 4.62 |
| Flye | 1,938,102,921 | 86,442 | 78,533 | 6,634 | 1.56 | 96.7 | 1.32 |
| Purgehaplotigs | 1,141,048,412 | 29,778 | 118,604 | 2,918 | 1.28 | 95.38 | 2.97 |
| RagTag | 1,141,383,012 | 26,432 | 136,910 | 2,517 | 30.86 | 95.71 | 2.64 |
| Lrscaff | 1,419,357,101 | 18,665 | 210,194 | 2,059 | 3,699.35 | 96.04 | 1.65 |
| Rails & cobler | 1,420,251,705 | 18,552 | 210,907 | 2,050 | 3,385.16 | 96.37 | 1.32 |
| RagTag | 1,420,292,605 | 18,143 | 211,737 | 2,037 | 3,387.41 | 96.70 | 0.99 |
| ArbitR | 1,420,280,939 | 18,136 | 211,815 | 2,034 | 3,387.48 | 96.70 | 0.99 |
| Rascaf | 1,420,308,595 | 18,007 | 216,668 | 1,977 | 3,389.41 | 96.70 | 1.32 |
| Purgehaplotigs | 1,195,861,358 | 14,952 | 225,090 | 1,609 | 3,540.36 | 95.71 | 1.65 |
| Ragtag | 1,195,981,958 | 13,746 | 249,518 | 1,400 | 3,549.64 | 96.04 | 1.32 |
| Blobtools | 1,195,981,958 | 13,746 | 249,518 | 1,400 | 3,549.64 | 96.04 | 1.32 |
| Ragtag | 1,172,699,129 | 6,835 | 297,634 | 952 | 3,338.35 | 96.04 | 1.65 |
| Pilon | 1,172,662,393 | 6,835 | 297,589 | 952 | 3,338.46 | 96.04 | 1.32 |

**Supplementary Table 3. Functional annotation statistics from different databases**

| Database | No. of terms linked to mRNA | No. of mRNA with term | No. of gene with term |
| --- | --- | --- | --- |
| InterPro | 13,274 | 9,370 | 6,796 |
| Gene ontology term | 11,726 | 5,793 | 4,172 |
| Pfam | 13,425 | 9,370 | 6,796 |

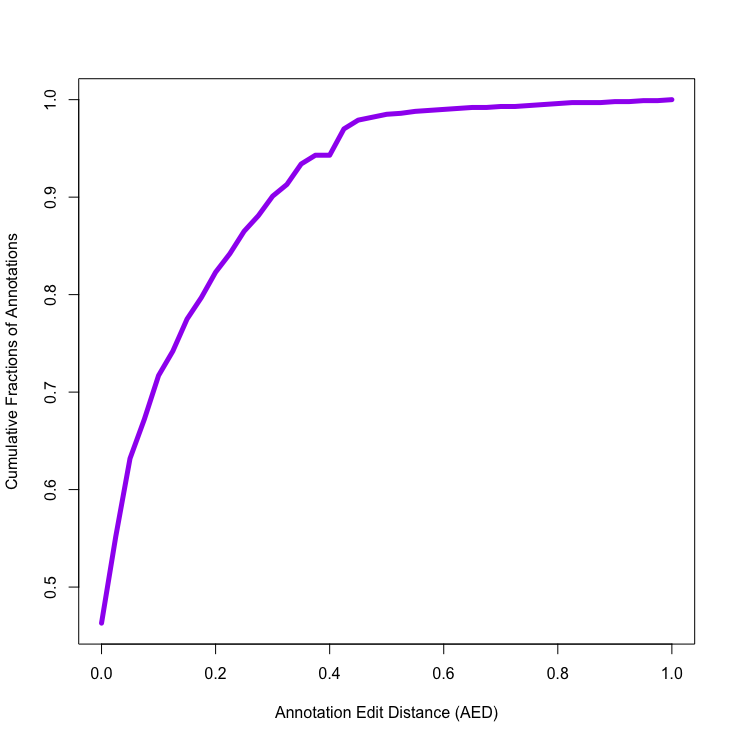

Supplementary Figure 1. Annotation Edit Distance of the predicted genes from three rounds of MAKER2 pipeline.
